# Combining biophysical models and machine learning to optimize implant geometry and stimulation protocol for intraneural electrodes

**DOI:** 10.1101/2023.02.24.529836

**Authors:** Simone Romeni, Elena Losanno, Elisabeth Koert, Luca Pierantoni, Ignacio Delgado-Martinez, Xavier Navarro, Silvestro Micera

## Abstract

**Objective:** Peripheral nerve interfaces have the potential to restore sensory, motor, and visceral functions. In particular, intraneural interfaces allow targeting deep neural structures with high selectivity, even if their performance strongly depends upon the implantation procedure and the subject’s anatomy. Currently, few alternatives exist for the determination of the target subject structural and functional anatomy, and statistical characterizations from cadaveric samples are limited because of their high cost. We propose an optimization workflow that can guide both the pre-surgical planning and the determination of maximally selective multisite stimulation protocols for implants consisting of several intraneural electrodes, and we characterize its performance in silico. We show that the availability of structural and functional information leads to very high performances and allows taking informed decisions on neuroprosthetic design.

**Approach:** We employ hybrid models (HMs) of neuromodulation in conjunction with a machine learning-based surrogate model to determine fiber activation under electrical stimulation, and two steps of optimization through particle swarm optimization (PSO) to optimize in silico implant geometry, implantation and stimulation protocols using morphological data from the human median nerve at a reduced computational cost.

**Main results:** Our method allows establishing the optimal geometry of multi-electrode transverse intra-fascicular multichannel electrode (TIME) implants, the optimal number of electrodes to implant, their optimal insertion, and a set of multipolar stimulation protocols that lead in silico to selective activation of all the muscles innervated by the human median nerve.

**Significance:** We show how to use effectively HMs for optimizing personalized neuroprostheses for motor function restoration. We provide in-silico evidences about the potential of multipolar stimulation to increase greatly selectivity. We also show that the knowledge of structural and functional anatomies of the target subject leads to very high selectivity and motivate the development of methods for their in vivo characterization.

## Introduction

Hybrid models (HMs) combining finite element modeling (FEM) and biophysical models of neural response [1] have been successfully exploited in the past to shed light on the main mechanisms of action of neural stimulation in the peripheral and central nervous systems [2]–[4]. HMs are powerful tools since they allow the prediction of the effect of stimulation at the level of single neurons, and rely on the same building blocks across different neuromodulation applications [1], [5]. Ideally, HMs can be used to evaluate and optimize different neural implants and stimulation protocols. In practice, the long computational time required to simulate a given experimental setup does not allow to explore the full space of the optimization variables in a continuous way, and thus HMs have only been used to provide proof-of-concept demonstrations and qualitative insights about a few previously chosen setups [2], [3]. Indeed, both FEM and biophysical models have important drawbacks which limit their large scale usability.

The optimization of a neuroprosthetic device requires defining the geometry of the electrodes, how they should be implanted in the target structure, and the stimulation protocol to produce the desired effects. In a peripheral neuroprosthesis, FEM can be used to compute from a candidate electrode geometry and implantation procedure a lead field matrix (LFM), which provides the sensitivity coefficients linking the currents injected by each of the stimulating sites in the implant to the corresponding extracellular potential along the nerve fibers. The response of the nerve fibers to the application of such extracellular potential is then typically computed using Hodgkin-Huxley-based multi-compartment models, the gold-standard being the McIntyre-Richardson-Grill (MRG) model of alpha motor fibers [6]. This allows to evaluate the effect of applying the given stimulation protocols from the implanted electrodes. To perform optimization, we need to repeat such evaluation for a potentially very high number of candidates, determined on the basis of the quality of the previously evaluated ones. The main objective of this work is to find a way to speed up the evaluation of the implant geometry, implantation, and stimulation protocols so that it is possible to perform their automatic optimization. Here, we propose to optimize implant geometry and implantation procedure using purely geometric objective functions, for example, maximizing the number of stimulation sites acting intra-fascicularly, or minimizing how close they are on average to the nerve fascicles. This allows to factor out the FEM from the implant geometry optimization, because its high computational cost would prevent the evaluation of many candidate geometries. The FEM is performed once the optimal implant geometry has been found, to obtain the LFM that is used to compute the extracellular potential applied to the fibers in the nerve.

As for the neural response computation problem, on the contrary, methods like the activating function formalism [7] and its further developments [8], [9] have been used in a few studies in the past [10]-[12] to enable the fast evaluation of a predetermined set of stimulation protocols. Nonetheless, these methods cannot be generalized to time-varying stimulation protocols, in which neural dynamics is strongly affected by the non-linear properties of the fibers, or to non-homogeneous or branched neural structures. Here, we propose to substitute them with machine learning-based surrogate models for the estimation of neural activation. They can be trained without specific expertise in biophysically accurate models, as they are black box models trying to infer the relation between the stimulation patterns applied to the neural units and their response computed through existing biophysical models and can be generalized to predict non-linear response properties. Specifically, we train multilayer perceptrons (MLPs) to predict the activation of MRG fibers from the pattern of applied extracellular potential and their diameter.

Thus, both the optimizations of implant geometry and stimulation protocols rely on complex black-box functions: the purely geometrical objective functions for the former, and the MLP surrogate model of neural response for the latter. To perform such optimizations, we chose to employ particle swarm optimization (PSO) [13], is a gradient-free evolutionary heuristics, in which a population of candidate solutions of an optimization problem is iteratively modified using the value of the objective function (also called “fitness”) evaluated at the candidate solutions and the information about their relative location in the space of optimization variables.

While the proposed framework can be generalized to several neuroprosthetic applications, here we apply our methods to movement restoration via intraneural stimulation of the human median nerve, following our recent experiments with non-human primates (NHPs) [14]. There, it was shown that the implant of a transverse intra-fascicular multichannel electrode (TIME) [15] in the median and radial nerves of a macaque monkey allows to produce the selective contraction of several muscles and even some functional grasps. Nonetheless, the development of the implanted electrodes and the determination of the implant site were performed on data from other individuals, and stimulation was applied through single site pulses with a-priori established amplitudes and pulse-widths. Here, we wanted to quantify which performances would be attainable if data from the target individual were available, and if multipolar optimized stimulation protocols were applied. Since the final aim of these studies is the translation in humans, we study here the problem directly using human data.

Our results also push towards the development of minimally invasive structural and functional imaging technique on peripheral nerve, since their availability would allow attaining results that are substantially superior to what has been shown in NHPs, notwithstanding the higher complexity of the targeted structures in humans.

## Methods

### Structure of the overall algorithm

The block diagram of the workflow proposed in the present article is shown in **Figure 1**. We will describe each step, referring to the corresponding figure panel and the section in the Methods where more details can be found.

**Figure 1.**
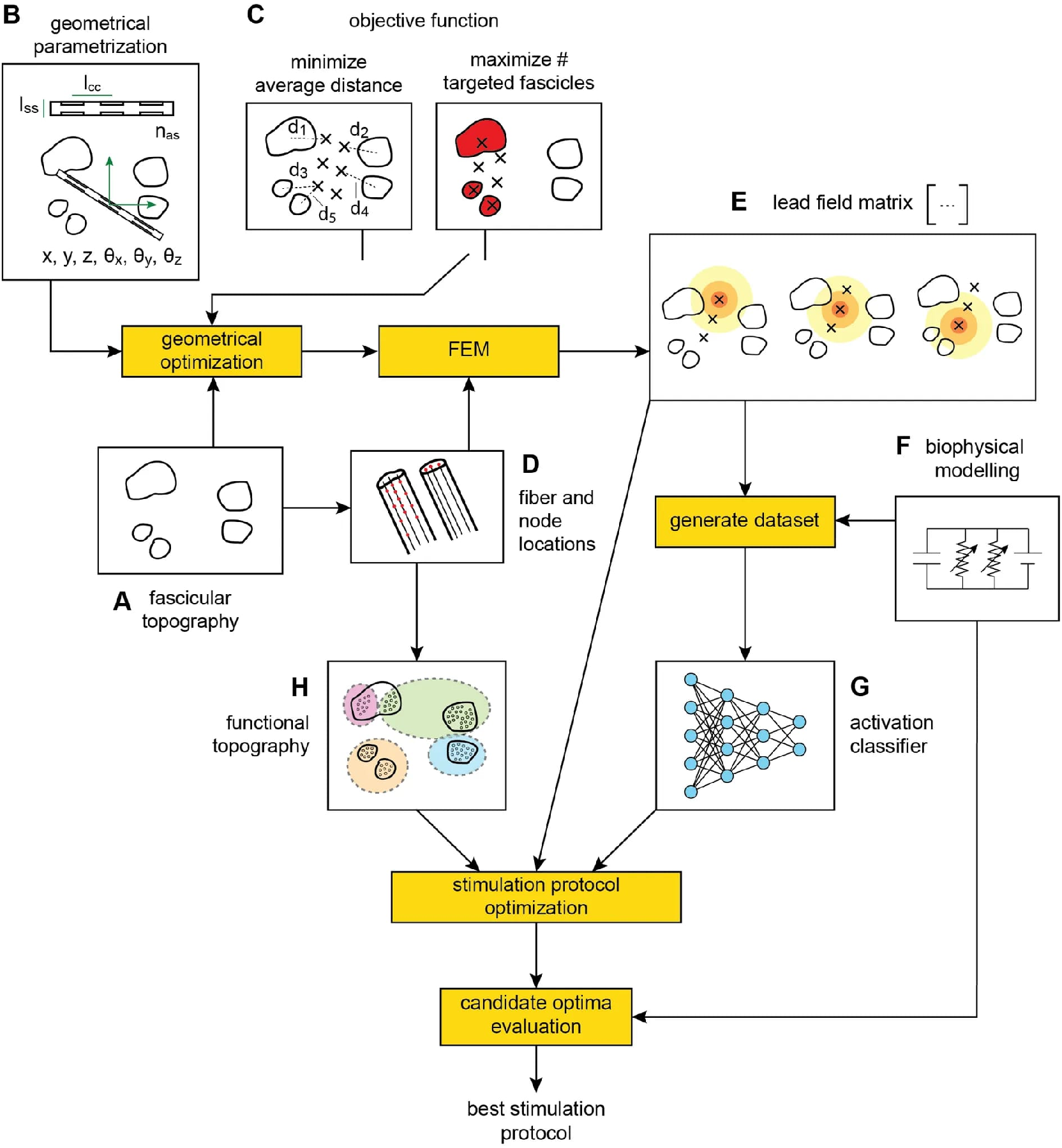
Proposed optimization workflow. (**A**) Parametrization of the electrode geometry and insertion. (**B**) Objective function for implant geometry optimization. (**C**) Fascicular topography of the nerve (fascicle boundary polylines in the nerve reference frame). **A**, **B**, and **C** are input to the geometrical optimization routine, which produces the optimal implant geometry input to the finite element modelling (FEM). (**D**) Determination of fiber locations in the nerve section and of fiber nodes along the nerve fiber path. (**E**) Characterization of the forward electrical problem by means of a lead field matrix. The optimal implant geometry and fiber node locations (**D**) are input to FEM to produce **E**. (**F**) Biophysical modelling of fiber stimulation. **E** and **F** are used to generate a dataset associating fiber stimulation and consequent activation. (**G**) Activation classifier trained on the generated fiber-stimulation-activation data. (**H**) Nerve functional topography. **E**, **G**, and **H** are input to stimulation protocol optimization, which produces a set of candidate optimal stimulation protocols. Candidate optimal stimulation protocols are evaluated using the biophysically detailed model (**F**) to establish the true optimal stimulation protocol.

The starting points are the fascicular topography of the target nerve (**Figure 1A** – see *“Fascicular topography”*) and a parametrization of the electrode geometry and insertion in the nerve (**Figure 1B** – see *“Geometrical parametrization of electrode geometry and insertion”*). These pieces of information are used to determine candidate-optimal implant geometries, according to a predetermined objective function linking the geometry of implants to their expected performance on the bases of heuristic considerations (**Figure 1C** – see *“Implant geometry objective functions”*). Putative fiber node locations are established (**Figure 1D** – see *“Fiber node location”*), and the LFM of the volume conduction problem is computed via FEM, using the nerve and implant geometries (**Figure 1E** – see *“Computation of the lead field matrix”*). A biophysically detailed model of the nerve fibers (**Figure 1F** – see *“Biophysical model of neural response”*) is employed in two points of our workflow: to produce a dataset of neural responses to different stimulation patterns that will be used to train a surrogate model of neural activation; and to “validate” candidate-optimal stimulation protocols. The surrogate model of neural activation is trained on the data produced employing the biophysically accurate model (**Figure 1G** – see *“Activation binary classifier”*). A probable functional topography is then established and used to convert fiber activations into fiber group recruitment and selectivity values (**Figure 1H** – see *“Activation, recruitment, selectivity”*), and used for the optimization of the stimulation protocols. The resulting candidate optimal stimulation protocols are then “validated” employing the biophysically detailed neural model.

### Median nerve fascicular and functional topographies

#### Fascicular topography

The “fascicular topography” of a nerve transverse section consists of the contours of the fascicles composing the nerve, and influences the outcomes of its stimulation as a thin, poorly conductive sheath (called perineurium) surrounds fascicles.

Human median nerve fascicular topographies have been obtained from the manual segmentation of stained histological sections, presented in Figure 5 of [16]. We considered the sections belonging to blocks X (here denoted as block 1), XI (here block 2), and XII (here block 3), which correspond to locations above the elbow, where fiber groups targeting all the muscles innervated by the median nerve are still present. In [16], each block was divided into three segments; nonetheless, this labelling of the nerve sections into segments was lost during the handling of the samples, and we needed to employ clustering techniques to recover it. We used the intersection over union (IoU) as a similarity measure between fascicular topographies, after aligning them using the iterative closest point (ICP) algorithm as implemented in the MATLAB function “pcregistericp”. For each cluster (corresponding ideally to the original segments), we determined the histological section with the highest average similarity with all other sections in the same cluster, and denoted it as the “representative histological section” for that segment. We then ordered the other sections in each segment with respect to their similarity to the representative section.

In **Supplementary Figure 1**, we show all the nerve fascicular topographies used in our simulations and the association between the different topography subsets employed to produce each different figure.

#### Simulated functional topography

With the term “functional topography” of a nerve section, we indicate the association map between the fibers populating it and their different targets (in this case, the different muscles that they innervate).

We defined 12 muscular groups, corresponding to the muscles innervated by the median nerve [16], namely: opponens pollicis (OP), abductor pollicis brevis (APB), flexor pollicis brevis (FPB), first lumbrical (1L), second lumbrical (2L), flexor digitorum superficialis (FDS), pronator quadratus (PQ), flexor digitorum profundus (FDP), flexor pollicis longus (FPL), palmaris longus (PL), flexor carpi radiali (FCR), pronator teres (PT).

Then, we followed the steps below:

- We divided the nerve into available regions, assigning zero available regions to very small fascicles (surface area < 0.05 mm^2^), one to small fascicles (surface area between 0.05 mm^2^ and 0.5 mm^2^), three to medium fascicles (surface area between 0.5 mm^2^ and 1 mm^2^), five to large fascicles (surface area larger than 1 mm^2^). We thus had

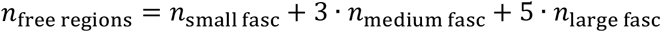
- We ordered the available regions with respect to distance from the centroid of the nerve section, computed as the centroid of all fascicle centroids.
- We removed a number of central regions, which were supposed to be occupied by sensory fibers. We arbitrarily chose to remove a number of regions equal to

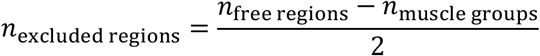
- For each muscle branch in [(OP, APB, FPB), (1L, 2L), FDS, (PQ, FDP, FPL), PL, FCR, PT]:

- If the branch contained only one muscle, we assigned its group to the free region closest to the center;
- Else, if there existed a fascicle containing a number of free regions higher than the number of muscles in the branch, we assigned it to that fascicle, else we randomly split the branch in two smaller branches and continued.

We filled the fascicles of each section with 25,000 uniformly distributed fibers following the axon counts presented in [16], and selected the motor fibers from this initial set of fibers according to the following steps. For each fascicle, fibers were assigned to the muscle groups in the fascicle one at a time. We selected the group to which we will assign a fiber by sampling a group identifier with a probability proportional to the number of fibers left to assign to each group. Then, we sampled the fiber to assign to the group by sampling a fiber with probability proportional to the value of the Gaussian density function assigned to the given muscle to each fiber.

In **Supplementary Figure 2**, we can see the different functional topographies and the corresponding optimal implants employed in the different simulations.

#### Fiber model and fiber node locations

The nerve fibers were modelled using the classic MRG model presented in [6] in its NEURON + Python implementation presented in [17]. Fiber diameters have been sampled according to a uniform distribution from the interval (12, 20) μm, which corresponds to the diameter limits for Aα motor fibers in the Erlanger-Gasser classification.

Fibers are divided into nodes of Ranvier and internodes, with internodes further divided into MYSA, FLUT and STIN. The center locations of each of these different sections (nodes of Ranvier, MYSA, FLUT, and STIN) are called “nodes” of the fiber. We modelled 40 internodes (and 41 nodes of Ranvier) for each fiber, and divided the corresponding nodes into four sets: a “lower boundary set”, a “FEM set”, a “propagation set”, and an “upper boundary set”. The FEM set is associated to a “FEM region”, whose length *ℓ*_FEM_ is set to 20 mm here. Starting from *z* = 0, we set a node of Ranvier of each fiber at a random location in (−*ℓ*_internode_/2, *ℓ*_internode_/2), where *ℓ*_internode_ is the length of the fiber internode (depending upon the fiber diameter). We then added Ranvier nodes for negative and positive z-values until we got outside the FEM region defined by the interval (–*ℓ*_FEM_/2, *ℓ*_FEM_/2). Five further nodes of Ranvier were added below the FEM region and the remaining nodes of the fiber were added above the FEM region. We recorded the time course of the membrane potential at the sixth node from the top extremity of the fiber, thus called “recording node”. The nodes below the FEM region constitute the lower boundary set, the ones above the recording node constitute the upper boundary set, and the ones between the FEM set and the upper boundary set constitute the propagation set. For a schematic representation of the subdivision of fiber nodes into the different simulation regions, see also **Supplementary Figure 3**.

In all stimulations, we applied a single current pulse of duration 5 ms. The amplitude of such current pulse coming from multiple stimulating sites (multipolar stimulation) is multiplied by the LFM to obtain the extracellular potential at the fiber nodes. The LFM had non-zero entries only corresponding to the nodes in the FEM set. To determine fiber activation, or, equivalently, whether an action potential is transmitted to the target muscles, we measured the membrane potential at the recording node, which is a large distance away from the region of the fiber where the stimulation is applied (the FEM region).

### Implant geometry optimization

#### Geometrical parametrization of electrode geometry and insertion

Here, we call “electrode” a single TIME, “implant” the set of TIMEs implanted in a given nerve section, and “electrode/stimulating site” each metal contact of the implanted TIMEs, through which the stimulation current is injected. We characterized each TIME through three geometrical parameters and six insertion parameters. The geometrical parameters were the number of active sites *n_as_* per arm, the arm-to-arm distance *ℓ_aa_* when the TIME is folded, and the center-to-center distance *ℓ_cc_* between consecutive active sites on a same arm of the TIME (also called “inter-site distance” in the following). The insertion parameters described the location (*x, y, z*) and orientation (*θ_x_, θ_y_, θ_z_*) of the electrode reference frame with respect to the nerve or laboratory reference frame. In the presented in silico experiments, we always assumed to optimize transverse electrode insertions (*θ_x_* = *θ_y_* = 0, *z* = 0) of electrodes having *ℓ_aa_* = 0.01 mm, and *n_as_* = 8 (corresponding to the electrodes employed in [18]). For a schematic interpretation of electrode and implant parameters, see also **Supplementary Figure 3**.

In the first in silico experiment (**Figures 2A-C**), we optimized both the insertion (*x, y, θ_z_*) for each implanted electrode and the inter-site distance *ℓ_cc_* common to all the electrodes in the implant. Next, we set *ℓ_cc_* = 0.75 mm [18] and optimized only the insertion parameters for each electrode. When testing two-electrode implants, we refer to the insertion variables for each electrode 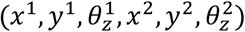 as an “insertion pair”. The optimization variables *x* and *y* were constrained to the interval (−5, 5) mm (slightly larger than the surface occupied by the nerve topography), *θ_z_* was constrained to (0, 2π), and *ℓ_cc_* to (0,1) mm.

**Figure 2.**
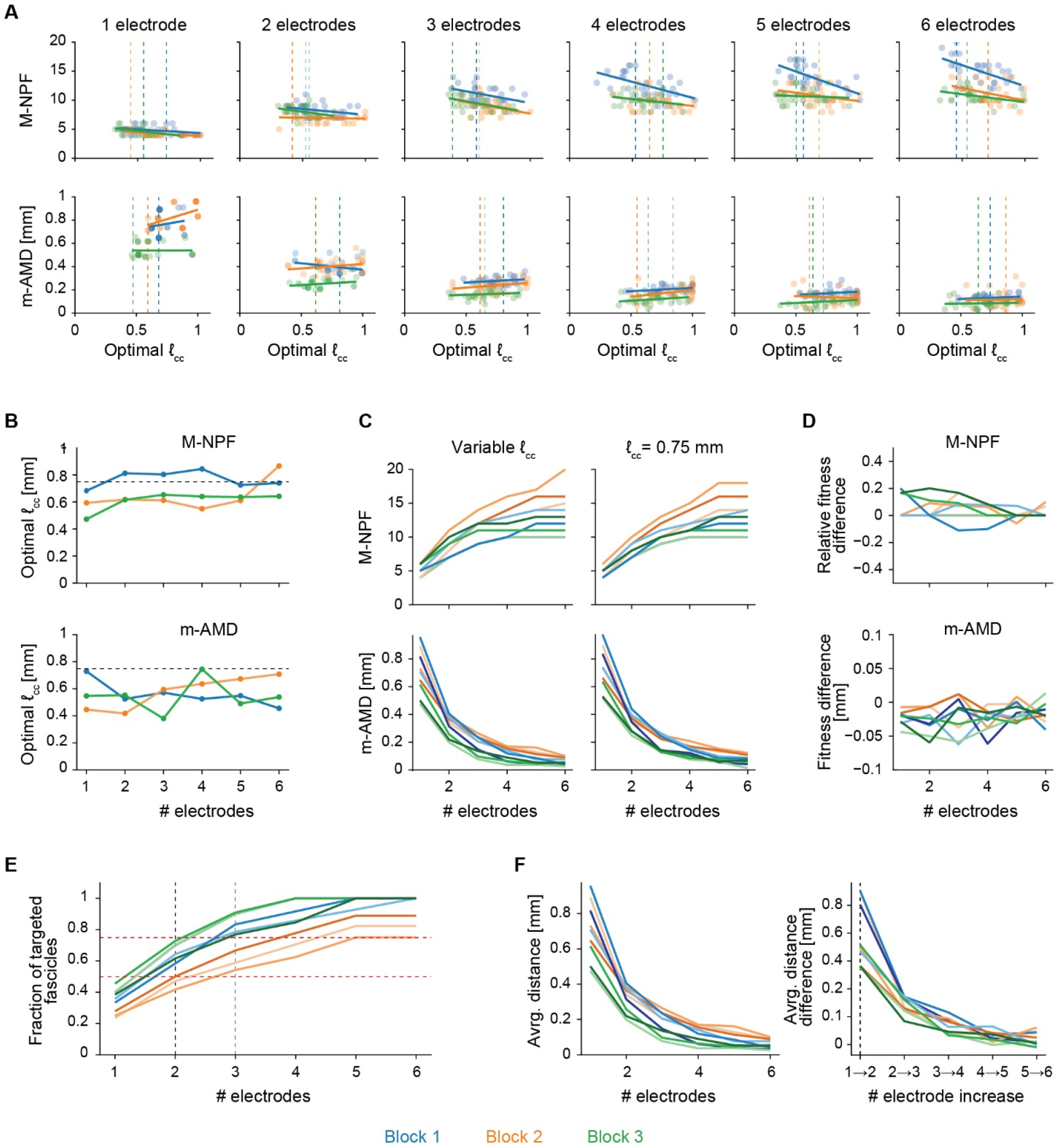
Dimensioning and choice of the number of implanted electrodes. (**A**) Relation between candidate optimal distance between active sites and geometrical fitness for different numbers of electrodes. Each point is a candidate optimal implant, the dotted lines correspond to the inter-site distance of the best-fitting geometry for each block, the dashed lines are obtained by linear regression of the points for each block. (**B**) Comparison between optimal inter-site distances found through optimization (colored solid lines) and TIME inter-site distance currently in use for human applications (black dotted line). (**C**) Fitness of the best geometries for different number of electrodes with variable or fixed inter-site distance. (**D**) Fitness difference (absolute for m-AMD, relative for M-NPF) between the best electrodes obtained with variable or with fixed inter-site distance. (**E**) Choice of the optimal number of implanted electrodes for M-NPF. (**F**) Choice of the optimal number of implanted electrodes for m-AMD, we show the fitness increments relative to adding the one implanted electrode at a time and then indicate with a vertical line the step leading to the highest fitness increment.

#### Objective functions

We tested two objective functions for implant geometry optimization: the maximization of the number of fascicles containing at least one stimulating site (M-NPF), or the minimization of the average distance between each fascicle and the closest electrode site (m-AMD). These objective functions can be used alternatively to maximize the number of sites acting intrafascicularly (shown as a desirable property of TIME implants [14]), or the “coverage” of the neural structure, respectively. It could also be possible to use composite measures obtained by linear combination of the two objective functions, but we did not perform such analyses here.

First, the maximization of the number of pierced fascicles (M-NPF), where the implant fitness corresponds to the number of nerve fascicles containing at least one implant stimulating-site

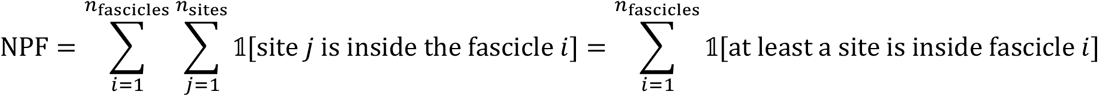

where *n_fascicles_* is the number of fascicles in the nerve section, *n_sites_* is the number of stimulating sites in the implant, and 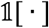 is the indicator function, which equals one if the argument proposition is true and zero otherwise.

Second, the minimization of the average minimum distance between fascicles and stimulating sites (m-AMD), where the cost of an implant is computed as the average of the distance between each fascicle and the closest stimulating site in the implant, or

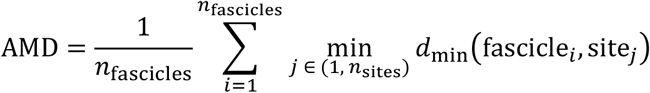

where *d*_mjn_(·, ·) indicates the minimum distance between two geometrical objects, or the distance between the closest points belonging to the two objects.

#### Optimization settings

We used the PSO algorithm implemented by the MATLAB function “particleswarm” with a swarm size of 50 and performed 150 iterations. The swarm size and number of iterations were chosen so that the fitness values generally converged before the last iteration. For all other parameters, we left the default values.

### Optimal insertion clustering

In order to evaluate the clustering of the optimal electrode insertions, we introduced a measure of distance between TIMEs. The computation of such distance measure is performed in two steps. First, the active sites of two TIMEs with the same number of active sites are coupled so that the distance between associated sites is minimized, or:

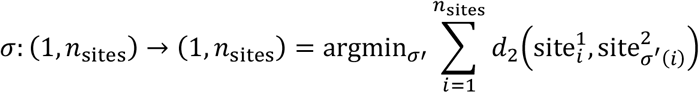

where the function *σ*(·) is a permutation function, *n*_sites_ is the number of stimulating sites in each TIME, and *d*_2_(·, ·) indicates the Euclidean distance between two points.

Then, the average of the distances between the associated sites is computed giving the desired distance between electrodes:

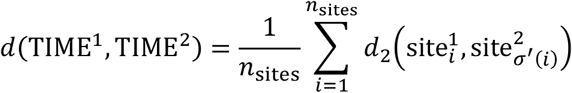

Using the above definition, we computed the pairwise distance between all the optimal electrode insertions and performed agglomerative clustering to highlight the similarity structures between insertions. The resulting dendrogram was cut where the curvature of the merging cost was the highest (elbow method).

### Computation of the lead field matrix

We built a geometrical model of the nerve extruding the poly-linear fascicular topography of a nerve section for a length of 20 mm. Point current sources were located in the implant site locations determined via implant geometry optimization. A saline bath with radius 10 mm whose upper and lower faces are located in the same planes as the nerve upper and lower faces surrounds the nerve and its external surface is grounded (*V* = 0).

One FEM solution is computed for each stimulating site in the implant, with a single stimulating site injecting a 1 μA current and all other sites switched off. The resulting potential map is interpolated using COMSOL default interpolation mechanism to obtain the LFM entries corresponding to fiber nodes in the FEM region. The LFM entries corresponding to the other fiber nodes were put to zero.

### Activation binary classifier

We trained a different MLP classifier for each implant geometry and for each number of concurrently stimulating active sites (one or three). All trained MLPs had the same topology (three layers with 32, 16, 4 units respectively) and were trained using the same hyper-parameter values (default values for MATLAB “patternnet”, which include the choices of hyperbolic tangent activation functions for the hidden layers and softmax activation function for the output layer).

We computed using FEM the LFM corresponding to the implanted electrodes and the 25,000 fibers randomly placed inside the section fascicles that we referred to in the section *“Simulated functional topography”*. To obtain each sample of the 10,000 constituting a training dataset, we considered one of the randomly placed fibers, we sampled randomly the identifiers of the stimulating sites (one or three identifiers, depending on the stimulation polarity) and the applied currents. We used the LFM to compute the extracellular potential applied to the fiber, and input it to our biophysically accurate models. A nerve fiber subject to a given stimulation protocol is considered active if at least a spike propagates to a sufficiently far away distance from the stimulation site. The presence of a spike is identified by checking for the presence of peaks higher than 20 mV with prominence 50 mV in the membrane potential time-course. In order to establish that we are recording the membrane potential time-course at a distance far enough from the stimulation site, we visually check that the membrane potential at the recording site is not strongly affected by the stimulation pulse. If this is the case, we can assume that the impulse will successfully propagate to its target.

We then randomly subsampled the most represented class (activated or not activated fibers) so that we had a balanced dataset, and trained a MLP using this dataset. To estimate the performance attainable by our classifiers, for 50 times we randomly partitioned the training dataset corresponding to each configuration into two subsets according to a 99/1 proportion, and computed the accuracy of classification attained on the smaller subset by a MLP trained on the larger subset.

### Stimulation protocol optimization

#### Stimulation protocol parametrization

For the stimulation protocol, the current amplitude of each active site was constrained in the range (—250, 250) μA for tripolar stimulation and (—250,0) μA for monopolar stimulation, unless stated otherwise. We chose a symmetric interval of amplitudes in the case of tripolar stimulation in order to allow current steering. When performing monopolar stimulation, negative current (cathodic) stimulation was chosen because it displays lower threshold values for fiber activation. The value of 250 μA was chosen as, in general, stimulation protocols attaining high selectivity values require low currents. In fact, the higher the injected currents, the higher the probability of recruiting more than one fascicle, thus lowering selectivity.

In order to optimize *n*-site stimulation without the need to decide beforehand which *n* sites among all available sites will be used for stimulation, we evolve a set of weights together with the stimulation amplitudes. Each weight refers to a single site, weights are constrained to vary in (0,1), and for each candidate solution the actual stimulation protocol is given by the amplitudes of the stimulating sites with the *n* highest values for the weights.

#### Objective function

The objective function for stimulation protocol optimization is the maximization of a selectivity measure that we define as

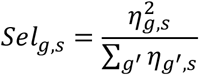

where *η_g,s_* denotes the value of the recruitment for group *g* and stimulation protocol s. We just remark here that such measure is obtained from the traditional selectivity measure introduced by Veraart and colleagues in [19]

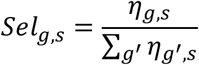

multiplying it by the recruitment of the target group.

The recruitment value *η_g,s_* corresponding to a fiber group *g* and a stimulation protocol *s* is computed as the fraction of active fibers in the group, or

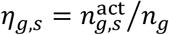

#### Optimization settings

We used the PSO algorithm implemented by the MATLAB function “particleswarm” with a swarm size of 25 and performed 50 iterations. The swarm size and number of iterations were chosen so that the fitness values generally converged before the last iteration. For all other parameters, we left the default values.

#### Evaluation of candidate optimal stimulation protocols

During stimulation protocol optimization, we employ the MLP classifiers to predict neural activations caused by candidate stimulation protocols, and PSO selects iteration after iteration the protocols leading to high selectivity for a given muscle group. It is possible for the PSO to exploit regions of the search space where our classifiers are not accurate and predict higher values of selectivity than the actual ones. Thus, once the PSO identified the five best candidate stimulation protocols, we computed the activation, recruitment and selectivity patterns values obtained via biophysically accurate models. Because we need to evaluate a relatively large pool of candidates, we only compute the response from a tenfold subsampling of the total fiber population in the nerve.

#### Effect of suboptimal electrode placement on optimal protocols

We analyzed what is the effect of a suboptimal electrode insertion on selectivity. First, we simulated imperfect electrode placement during surgery by slightly perturbing the optimal insertion parameters *x, y, θ_z_* by Δ*x*, Δ*y* sampled uniformly in (−0.25, 0.25) mm, and Δ*θ_z_* sampled uniformly in (−15°, 15°) for each of the two implanted electrodes. Second, we simulated imperfectly predicted fascicular topography by applying to the representative section of block 1 the optimal implants obtained for two different sections of the same block, having an intersection over union (IoU) with the reference section of 0.8 and 0.46, respectively. Third, we simulated a random electrode insertion by randomly sampling *x, y, θ_z_* with the constraint that no active site was outside a square of side 6 mm centered at the origin.

## Results

### Optimization of implanted electrode design

First, we show how our geometry optimization routine can be used to provide guidelines to define the optimal size of the electrode to implant in a given nerve. In **Figure 2A**, we can observe that the candidate best inter-site distances fall into a wide interval (from 0.5 mm to 1 mm approximately), but that lower inter-site distances lead in general to slightly better performance using both the objective functions. In **Figure 2B**, we notice that there is no evident relation between the best inter-site distances and the implanted nerve block or the number of electrodes constituting the implant.

At the same time, our geometry optimization routine allows to evaluate the performance of existing electrode designs with respect to the optimal one. To show this, we ran the insertion optimization for implants composed of electrodes with a fixed inter-site distance equal to 0.75 mm, value that corresponds to the electrode design employed by our group in the latest clinical trials to restore hand sensation in trans-radial amputees [18]. In **Figure 2B**, we can see that the optimal inter-site distance is in general lower than the one of the existing designs. Nonetheless, in **Figure 2C**, we directly compared the performance attainable by the fixed and the optimized design, showing that the loss in performance that derives from using our fixed design is not substantial.

### Establishing the best number of implanted electrodes

Our geometry optimization routine can be also used (through the analysis of the performance curves in **Figure 2C**) to establish how many electrodes should be implanted at the different levels along the longitudinal course of the nerve to obtain the highest performance with the lowest invasiveness. **Figure 2D** shows that two electrodes optimized according to M-NPF allow targeting more than 50% of the fascicles for the two most proximal nerve blocks, and three electrodes target more than 50% of the fascicles for all blocks and more than 75% of the fascicles for the two most proximal blocks. In the case of m-AMD optimization, we can see that increasing from one to two electrodes grants the maximal fitness increase with respect to adding one electrode to other configurations, and thus two electrodes should be preferred as they grant the maximum fitness increase (**Figure 2D**). In the following, we will always consider the implant of two electrodes, whose feasibility has been already shown in the human median nerve [18], [20].

### Characterization of the best strategies for electrode insertion

In this section, we show how candidate implant insertions can be analyzed. In **Figure 3A**, we see the candidate optimal electrodes obtained performing 20 independent PSO runs on the same nerve section. We can see that many electrodes in different implants occupy very similar locations in the nerve section, suggesting that there may be a low number of particularly advantageous single-electrode insertion strategies. We thus introduced a measure of dissimilarity between TIMEs and used it to perform hierarchical clustering (see **Methods**, *“Optimal insertion clustering”*) to highlight the similarity structures between the proposed insertions. **Figure 3B** shows the obtained dendrogram and its cutting; the resulting clusters of electrode insertions are shown in **Figure 3C**. In general, different clusters contain different numbers of insertions and display different extents of variability, which can be seen as the robustness of the optimal insertion strategies to variations of the insertion parameters.

**Figure 3.**
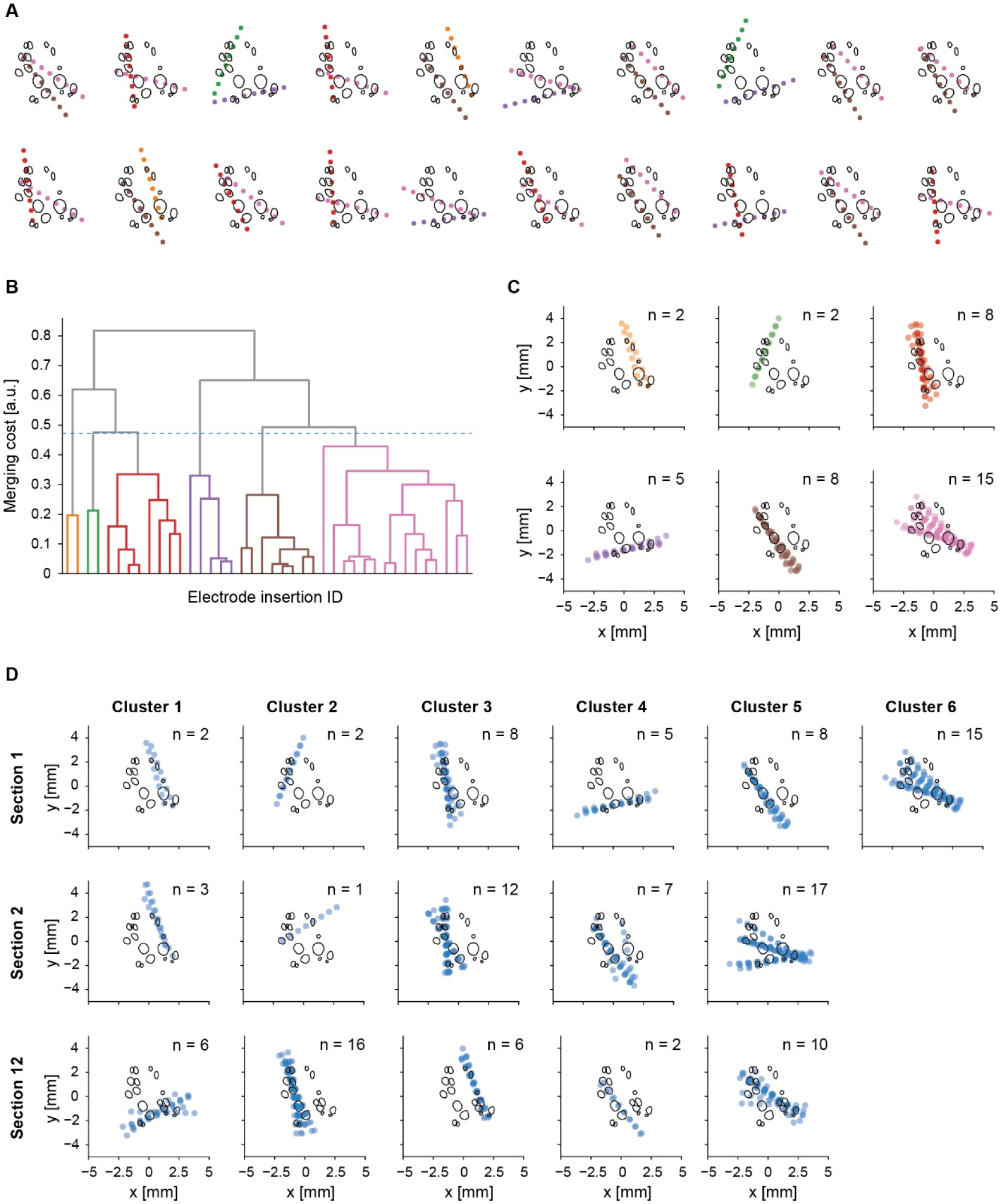
Multiple insertion optimization routines on the same section. (**A**) Optimal implant insertions for a single nerve section; each subplot corresponds to a random initialization of the optimization algorithm (20 in total). (**B**) Dendrogram for the clustering of electrode insertions; the dashed line corresponds to the level of the cut. (**C**) All electrode insertions from the clusters corresponding to plots A and B. Cluster colors are coherent among panels A, B and C. (**D**) All electrode insertions divided into clusters obtained for the three sections; n indicates the number of electrodes belonging to that cluster (20 for each section).

The couples of electrode insertions occurring most frequently together are shown in **Table 1**. Only six of the thirty possible cluster couplings occur, and most of the optimal implants are constructed using electrode insertions from a few clusters. These insertions should be preferred when performing the surgical implantation. In **Figure 3D**, we show the clusters of electrode insertions corresponding to different sections in the same nerve segment. We can see that insertion clusters are robust with respect to small variations in nerve topography.

**Table 1.** Frequency of occurrence of insertion pairs. Number of candidate optimal implants (out of 20 repetitions) that contain one electrode inserted according to cluster A and one electrode inserted according to cluster B from **Figure 2A-C**. In bold, the two most frequent cluster pairs.

| Insertion A | Insertion B | # of occurrences |
| --- | --- | --- |
| 1 | 5 | 2 |
| 2 | 4 | 2 |
| 3 | 4 | 1 |
| **3** | **6** | **7** |
| 4 | 6 | 2 |
| **5** | **6** | **6** |

### Training and accuracy of the activation classifier

For each experimental setup (choice of nerve level and implant geometry), a MLP binary classifier was trained to predict the fiber activations produced by biophysically accurate models in that specific setup. The expected classification accuracies were computed performing Monte Carlo cross validation for 50 iterations with a 99%-1% split of the available labelled datasets. Further details on the classifiers and the training process can be found in **Methods**, *“Activation binary classifier”*. In **Table 2**, we report the obtained estimated accuracies and the number of samples in the balanced labelled dataset used as the base for Monte Carlo cross-validation and for the subsequent final MLP training. Accuracy always exceeded 0.95 except for the single case of block 2 using m-AMD in monopolar stimulation, for which accuracy was equal to 0.94.

**Table 2.**
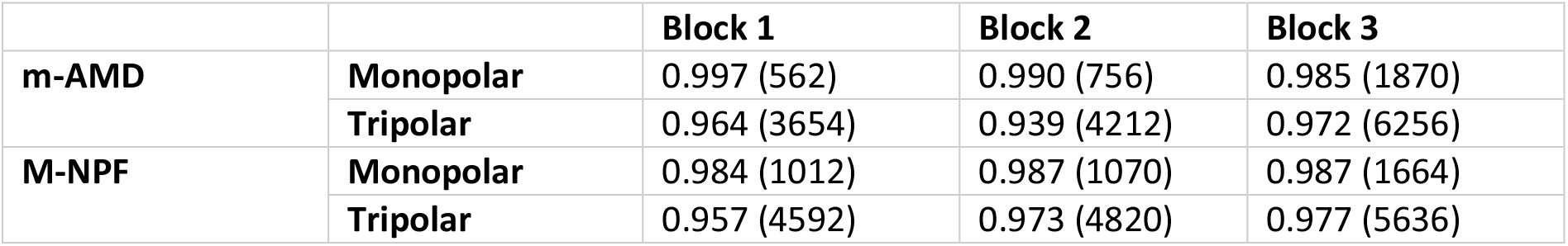
Activation binary classifier estimated accuracy and training set size. The estimated accuracy of each classifier is obtained performing Monte Carlo cross-validation, 50 samplings with a training/test set proportion of 99%/1%. Between parentheses, the number of samples constituting the largest balanced dataset obtained from 10,000 random stimulation protocols, which was used to perform the training of the final MLP. One classifier is trained for each combination of geometric objective function, number of stimulation sites, and nerve fascicular topography.

Even though the obtained accuracies are very high, it could happen that the optimization routine generates adversarial examples for these classifiers, predicting selectivity values much higher than the actual ones. In **Methods**, *“Evaluation of candidate optimal stimulation protocols”*, we show that in general this is not the case. In the following, we “validate” the candidate optimal stimulation protocols by computing their selectivity on a ten-fold subsampling of the fiber population, and report the results corresponding to the most selective protocol.

In **Figure 4A**, we plot the group-wise selectivity values obtained using the three methods. For the MLP classifier and under-sampled estimated selectivity, we plot the best selectivity across five independent re-initializations of the algorithm. We then evaluate the protocol attaining the best selectivity using the full population. We can notice that undersampling the fiber population leads to a systematic overestimation of the selectivity. To understand better this phenomenon, we investigated the effect of under-sampling on recruitment. In **Figure 4B**, we can see that under-sampling tends to overestimate high recruitment values and to underestimate low recruitment values, which results to a systematic overestimation of the performance of high selectivity stimulation protocols (where the recruitment of the target group is overestimated, and the recruitment of the non-target groups is underestimated). Nonetheless, the average error on recruitment is very low for both the under-sampled model and the MLP classifier. In **Figure 4C**, we display the recruitment patterns estimated using the three methods, corresponding to the highest estimated selectivity for each muscle group.

**Figure 4.**
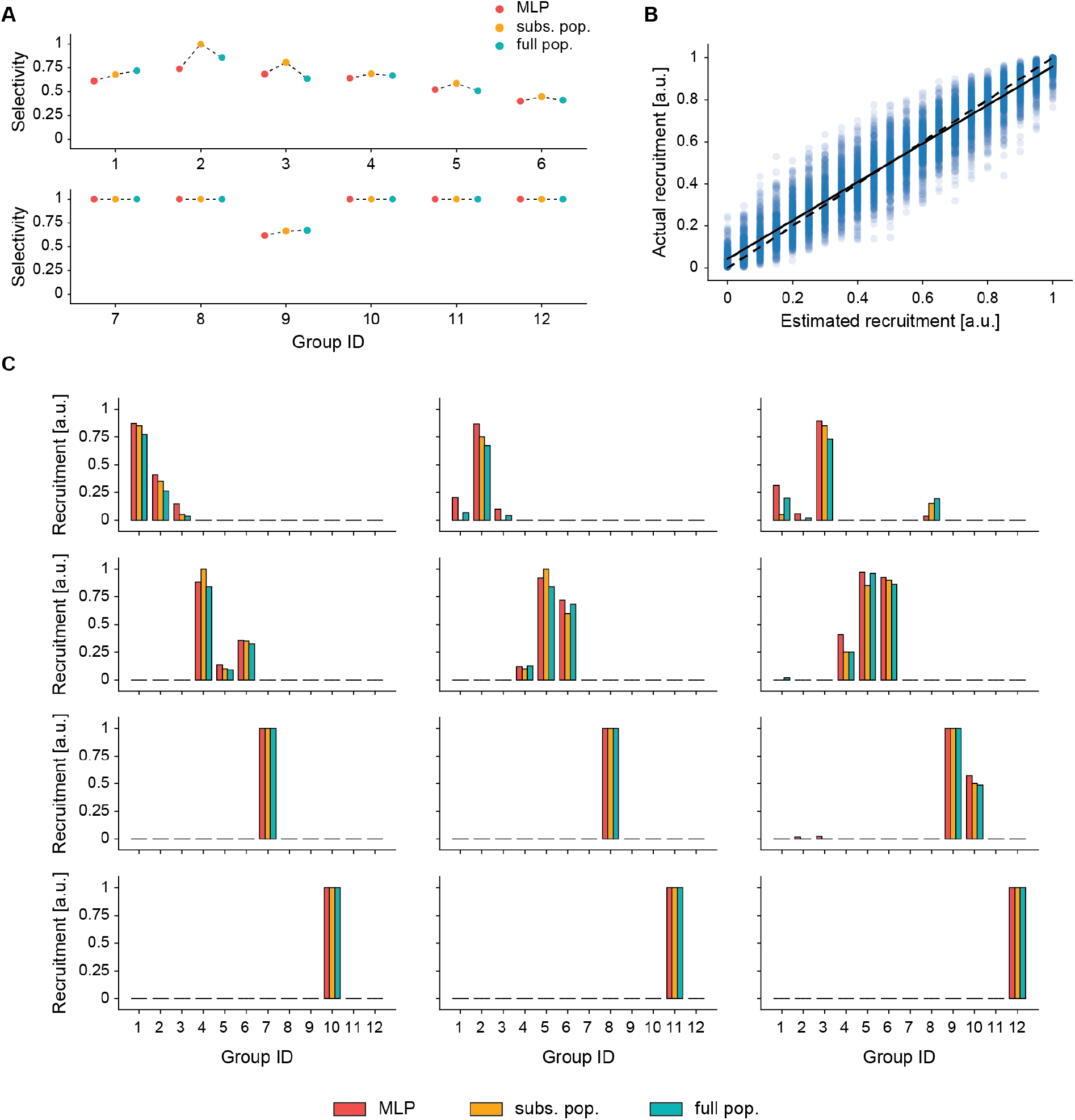
Candidate optimal stimulation protocol evaluation. (**A**) Group-wise selectivity values corresponding to candidate optimal stimulation protocols. For each group we provide: selectivity computed using our activation classifier, a subsampled fiber population, or the full population. (**B**) Effect of subsampling on recruitment. The true recruitment is compared with the recruitment estimated from the subsampled population. The dashed line is a linear regression on all points, and the solid line is the quadrant bisector, provided for comparison purposes. (**C**) Recruitment patterns for the best protocols evaluated with our activation classifier, fiber subsampling, and the full fiber population.

### Optimal stimulation protocols in different conditions

We performed several in silico experiments to characterize the performance of the optimal stimulation protocols found by our routine in different conditions.

First, we compared monopolar and tripolar optimal stimulation protocols. In **Figure 5A**, it can be seen that monopolar stimulation is substantially less effective than tripolar stimulation. The total selectivity for monopolar stimulation is 0.45 and for tripolar stimulation is 0.78. Comparing the results from **Figure 5A** with the group locations in **Supplementary Figure 2**, we can see that muscle groups that do not show substantially different selectivity for monopolar and tripolar stimulation are generally in single-group fascicles having a close by electrode site, which allows selective monopolar stimulation. Instead, when the target muscle group shares its fascicle with other muscle groups or when such fascicle does not have a stimulating site close by, current steering is required to improve the stimulation selectivity. In the rest of our work, we focus on the optimization of tripolar stimulation.

**Figure 5.**
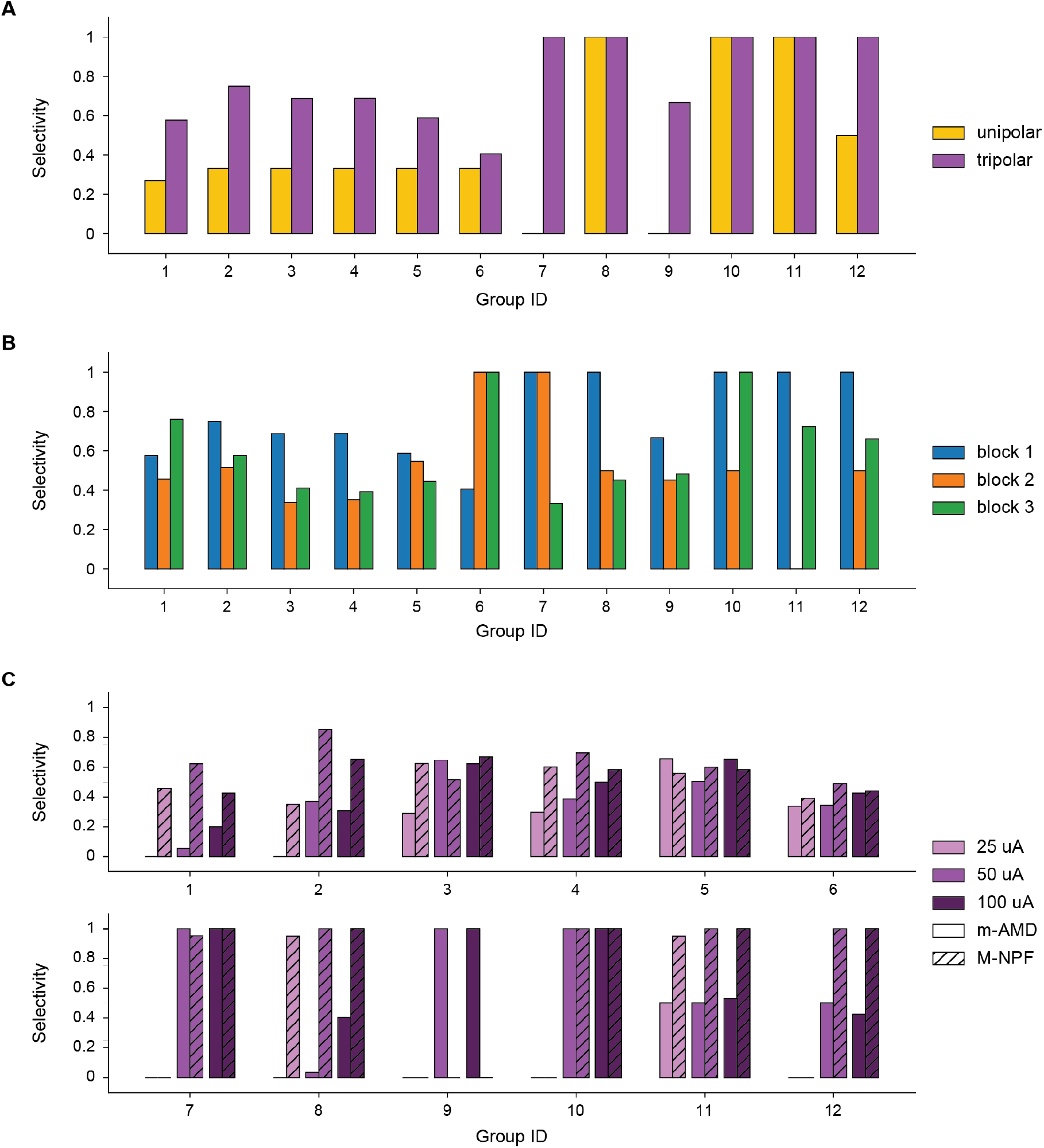
Stimulation protocol optimization in different conditions. (**A**) Comparison between monopolar and tripolar stimulation, optimization on the reference section from block 1. (**B**) Comparison between tripolar optimal stimulation for the reference section from blocks 1, 2, and 3. (**C**) Comparison between the best tripolar stimulation for m-AMD and M-NPF optimized implants, when imposing different maximal site-wise current values. All values refer to the reference section from block 1. All selectivity values correspond to the selectivity measure introduced in the present work.

Second, we compared the selectivity attainable when implanting and stimulating at different levels along the course of the nerve to establish, together with surgical accessibility considerations, the best implantation location. We observe in **Figure 5B** that the group-wise selectivity has generally the highest values in the most distal block (block 1), followed by the most proximal block (block 3), while the lowest values correspond to the middle block (block 2). Coherently, the total selectivity values obtained were 0.78 for block 1, 0.60 for block 3, and 0.51 for block 2.

Third, we compared the performance of the optimal implants according to the M-NPF and m-AMD fitness functions. We performed stimulation protocol optimizations constraining the current amplitude in different intervals to compare intrafascicular (M-NPF) and intraneural (m-AMD) optimization strategies at different current levels. The results are shown in **Figure 5C**. For currents in (−50, 50) μA, we had m-AMD = 0.17 and M-NPF = 0.40, for (−100, 100) μA we had m-AMD = 0.53 and M-NPF = 0.73, for (−200, 200) μA we had m-AMD = 0.59 and M-NPF = 0.70. m-AMD generally performs worse than M-NPF and while its performance is particularly bad for very low current values, M-NPF is more robust.

Finally, we analyzed what is the effect of a suboptimal electrode insertion on selectivity (see **Methods**, *“Effect of suboptimal electrode placement on optimal protocols”*). In the case of simulated imperfect electrode placement during surgery, we obtained a total selectivity of 0.63. In the case of imperfectly predicted fascicular topography, the total selectivity values obtained after optimizing the stimulation protocol were 0.59 and 0.51, respectively. In the case of random electrode insertion, we obtained a total selectivity value of 0.35.

## Discussion

We have presented a pipeline for the optimization of the design, implantation, and use of implantable neuroprosthetic interfaces (**Figure 1**). The implant geometry is optimized automatically through a geometrical fitness function that exploits anatomical data of the subject to implant and is then used to train a ML predictor of fiber activation that allows performing stimulation protocol optimization. While the steps of our pipeline are general, all our experiments were performed using TIMEs, and median nerve topographies, with a movement restoration application in mind. After presenting our results, we will also discuss what difficulties can arise when we try to generalize the presented workflow to other electrodes, nerves, and applications.

In the present work, we separated the problems of finding an optimal implant geometry and insertion, and of finding the optimal stimulation protocol for the optimal implant. Ideally, implant geometry, insertion, and stimulation protocols should be optimized jointly, as the specific implant geometry and insertion affects which stimulation protocols are best and what selectivity they produce. Nonetheless, changing implant geometry or insertion requires to update the LFM computed through FEM and then used to compute the effect of the applied stimulation protocols. This is not possible because no surrogate model of FEM is currently available. This is also the reason why we needed to use heuristic, purely geometric objective functions to optimize implant geometry and insertion. The chosen objective functions agree with our intuition of how selective electrical stimulation works. The M-NPF objective function has been defined noticing that stimulating intra-fascicularly generally increases selectivity (see also [14]). This is reasonable, as fascicles are surrounded by a poorly conductive sheath and should contain, at least distally, one or a few muscle groups, helping to target them selectively. Instead, the m-AMD objective function encourages having at least one active site very close to each nerve fascicle, thus being able to stimulate all fascicles with relatively low intensity.

We have shown how to employ our routine to find the optimal geometry of a set of intraneural electrodes for peripheral nerve stimulation (**Figure 2A-B**), and to evaluate the expected performance of an existing electrode geometry when optimally inserted (**Figure 2C**). This can help the experimenter to evaluate the opportunity of employing an existing electrode geometry instead of asking the manufacturer for a personalized geometry.

The optimal number of electrodes to be implanted has been established by analyzing the trade-off between invasiveness and geometrical fitness (**Figure 2D**). We could not optimize the electrode count directly in the PSO as the optimization would have been biased towards high values, which allow a better coverage of the neural structure. Moving towards automatic optimization of neuroprostheses requires to identify and avoid similar biases in the optimization protocols.

The problem of where the implantation should be performed along the longitudinal axis of the nerve has been discussed. In **Figures 2C-D** we can see that the number of fascicles targeted by the optimal implants is higher in more proximal locations, while the proportion of targeted fascicles is higher in more distal locations. This poses a trade-off that can be solved by evaluating the optimal stimulation protocols. Using this criterion, we found that the best implantation site was the most distal section, followed by the most proximal, and finally by the central section (**Figure 5B**). A similar analysis has been presented by Badi, Wurth and colleagues [14], showing that in the macaque median nerve, too proximal and too distal implant locations lead to lower performances. The higher selectivity that we attained distally is likely due to the implantation of twice the number of active sites in optimized arrangements, while in [14] the insertion was manually set. The improvement in performance obtained proximally could be derived from the use of multipolar stimulation protocols, which allow coping with multi-group fascicles thanks to electric field steering (**Figure 5A**).

The output of our optimization routine for different random search initializations can be clustered in a low number of optimal electrode insertions (**Figure 3**), and the intra-cluster variability suggests how sensitive that optimal insertion strategy is to geometrical variations (**Figure 3C-D**). When we analyzed pairs of electrode insertions, we found that the number of represented implants is actually even lower, with only a couple of preferred configurations (**Table 1**). Moreover, the candidate optimal electrode insertions for similar nerve sections are in general very similar (**Figure 3D**). On one side, the fact that the number of optimal insertion strategies is low and replicable provides good implant guidelines to the surgeon and paves the way for automated implantation of peripheral nerves. On the other hand, the presence of a few alternative implantation procedures allows the surgeon to choose which one is more viable in the practice. We also quantified the loss in selectivity given by different suboptimal implant insertions, due to lack of precision in the determination of the nerve topography and in the implantation procedure, showing that random implant insertion leads to reduced selectivity.

The results of stimulation protocol optimization were provided in terms of a new selectivity measure, derived from the traditional one introduced by Veraart and colleagues in [19]. The traditional measure has the issue that low stimulation levels, can produce to very low recruitments of a single group without co-activation of other groups, which produces maximum selectivity while being non-functional. Multiplying the selectivity value by the target group selectivity, we solve such problem, making the selectivity a continuous function of group-wise recruitments. Our measure is always smaller than the traditional one, thus leading to more conservative results. In **Figures 4A-C**, our selectivity values can be compared to the corresponding recruitment patterns to gain some quantitative insight on our selectivity measure.

We show that multipolar stimulation provides substantially higher selectivity than unipolar stimulation (**Figure 4A**). This is an expected result, as the former allows more freedom to shape the generated electric potential field. Further investigations of multisite stimulation protocols should be carried out, to study the impact of the number of independently controllable stimulating sites on the maximal attainable selectivity. Such analyses may help to understand whether substantial technological advancements are needed to solve the problems related to increasing the number of independently controllable stimulating sites.

### Limitations and future developments

#### Assumptions on nerve structural topography

We obtained the 3D model of the nerve by extruding a single 2D transverse section. Such simple model cannot represent accurately the structure of nerves whose fascicular topography changes frequently along its path. While it is known that in the median nerve there are long tracts with very limited internal branching [21], in other nerves, like the vagus nerve branching is much more frequent [22]. Even in such cases, because neural activation is influenced by the extracellular potential applied to a relatively short fiber length around the stimulation level [23], it is possible that if individual nerve fibers do not displace substantially in the nerve transverse section, their activation thresholds would be approximated well enough. In any case, while there is no guarantee that the quality of our results may translate to more complex nerves, the presented optimization framework can be used as is, simply changing the nerve geometry model built in the FEM software and interpolating the extracellular potential at the locations of the fiber nodes in 3D, like shown in [24].

A more severe limitation of our method is that it requires some knowledge of the fascicular topography of the target nerve before surgery, in order to run our implant geometry localization routine. While it has been shown that it is possible to determine in a non-invasive way the fascicular topography of a nerve using 7T magnetic resonance [25] and ultra-high frequency ultrasound [26] imaging, the precision and reliability of these techniques have never been thoroughly quantified. The present study shows that such imaging techniques should be actively developed and characterized, as they can allow pre-surgical planning which leads to a substantial increase in the performance of the implanted devices with respect to the state of the art.

#### Assumptions on nerve functional topography

We followed these principles: (1) functional groups (sensory or motor) that branch out of the nerve more distally occupy more central locations in the nerve transverse section; (2) fibers innervating neighboring muscles should be closer in the nerve transverse section [16], [21], [27]. In the light of the latter principle, we further assumed that: the fibers belonging to a single muscle group are all in a single fascicle; fibers groups with similar functions should be preferably placed in a same fascicle. We identified three such macro-groups: OP, APB, and FPB (constituting the thenar eminence, innervated by the recurrent branch of the nerve); 1L and 2L (lumbricals, innervated by the palmar digital branch of the nerve); PQ, FDP, and FPL (innervated by the anterior interosseous branch of the nerve) [16]. Finally, since the sensory fibers going to the hand are the vast majority of the fibers populating the median nerve [16], and because of principle (1), we decided to exclude muscle fibers groups from a number of central regions, ideally dedicated to sensory fibers. We highlight that while we tried to follow all qualitative insight found in literature to generate the test functional topographies, such topographies should not be considered as an outcome of the present work, but rather as a piece of data that was required to compute the expected selectivity obtained after optimization. Our attention in trying to replicate a natural functional topography should lead to selectivity values that will translate well to future experimental settings.

When optimizing the stimulation protocols, we assumed to know the functional topography of the implanted nerve section. Intuitively, the approximate location of functional groups can be determined by applying stimuli from single sites located at different positions in the implanted electrodes and using triangulation (similarly to what is done using neural recordings in [28]-[30]). However, to the best of our knowledge, rigorous methods have not been developed yet. We have shown that the availability of a precise functional topography of the nerve allows attaining extremely good selectivity patterns, thus encouraging the investigation of functional localization methods.

While it would be ideal for development of fully personalized neuroprosthetic devices to have methods for the in vivo quantification of the nerve structures, their characterization via histological and immuno-histochemical analyses of cadaveric samples may provide valuable general guidelines. Since such analyses require adequate instruments, expertise, and large economic investments, initiatives like the SPARC data research center [31] are extremely relevant as they constitute a framework that allows the dissemination of such costly data collections throughout the scientific community.

#### Validation of the present approach

The fact that such methods of in vivo quantification of structural and functional topography are not available prevents the experimental validation of such results. While HMs in general are a validated technique [32], [33] there are many aspects of an experimental setting that could result in a much lower selectivity than the one found in our in silico set-ups. For example, the mismatch between the guessed and actual fascicular and functional nerve topographies, or the fact that the effect of multiple concurrently stimulating electrode sites may not superpose linearly. Further in silico and animal experiments could be performed before transitioning to human subjects, to adapt the presented methods so that the impact of these phenomena will not hinder the benefit of automatic model-based optimization.

#### Further limitations related to the implant

Finally, we ignored several issues that may occur after implantation. A first problem is related to the value of the electrode geometrical variable *ℓ_aa_*, which here we assumed constant since it corresponds to the thickness of the folded polyimide layer constituting the electrode body and thus cannot be easily modified. Nonetheless, during/after insertion the electrode arms can detach, leading to an additional separation between them, which was ignored here, but further studies should provide reliable estimates of its magnitude and investigate its impact on the stimulation outcome. A second problem is related to the foreign body reaction to the implant and stimulation. Such changes range from the swelling of the pierced fascicles [36], to the variation in time of the stimulation threshold due to the encapsulation tissue that surrounds stimulating sites [23]. These phenomena are known to substantially affect stimulation outcomes, but their characterization is far from complete and thus future studies will need to include them in computational frameworks like the present one.

#### Generalizability of our methods

Our geometric objective functions have been used to optimize some of the parameters of a set of implanted TIMEs. Nonetheless, they could be used as is also for other electrode families. For example, the M-NPF could be used to optimize the number of microelectrodes in a Utah array and their arrangement, and the m-AMD could be used to optimize the number of circumferential stimulating sites in a cuff electrode. New, implant-specific geometric objective functions can be introduced using the intuition behind selective stimulation with different electrode families. Indeed, the fact that our objective functions can be used out-of-the-box is not enough to guarantee that the corresponding results will be as good as the ones presented here. Further in silico analyses should be performed.

The same can be said for applications that target other parts of the nervous system. The basic building blocks of our approach can in fact be used to optimize spinal cord, brain, and retinal prostheses. As an example, the m-AMD objective function could be used to fix the geometry of an epidural spinal cord electrode array, where it is desirable to have stimulating sites close to each spinal root to be able to produce selective stimulation protocols.

To limit the complexity of our analyses, we limited the number of parameters to be optimized for each TIME, but we could have included in the search space for the optimization the electrode-wise variables *z, θ_x_, θ_y_* without any modification to what has been presented. Instead, the value of *n_as_* needs to be optimized by performing separate geometry optimizations with the different reasonable values for the parameter, as we have done to set the number of electrodes to be implanted within a single nerve, as higher stimulating site counts naturally tend to produce higher fitness values. The fitness values obtained for the best implants corresponding to each value of *n_as_* are then compared and a decision on the value to be employed is taken, keeping into consideration the complexity of building an electrode with such features, and the added invasiveness of the resulting interface.

All the steps following the computation of the LFM do not depend on the specific nerve or implant geometry and can be applied as is to electrode families and nerves different to the ones shown here. Depending on the target application, the stimulation protocol objective function may need to be modified or adjusted. Here, we optimized stimulation protocols to maximize selectivity, which is appropriate for movement restoration and bioelectronic medicine applications (where each functional group corresponds to a different innervated muscle and to a bodily function or innervated organ, respectively). Instead, sensory restoration, in which fibers account for sensations that span a continuum, poses a very different challenge, and thus an adequate objective function for stimulation protocol optimization should be determined and validated.

#### Joint optimization of implant geometry and stimulation protocols

In the present work, we optimized implant geometry and stimulation protocols in two sequential steps. Since the shape and the insertion of the implant obviously influence the optimal stimulation protocols and their selectivity, joint optimization of implant geometry and stimulation protocols will produce selectivity values higher than the ones presented here. Still, such joint optimization would require the computation of a new LFM each time that the geometry of the implant is modified, leading to extremely long objective function evaluation times, jeopardizing the possibility of performing optimization. The development of surrogate models of the FEM to evaluate the LFM corresponding to a given stimulation setup at a reduced computational cost will be required to introduce routines for the joint optimization of implant geometry and stimulation protocols.

## Conclusions

Our guidelines and methods provide a quantitative perspective on the advantages of automatic optimization pipelines for personalized neuroprosthetic devices, moving the role of modeling from qualitative assessment of different stimulation settings to being a fundamental tool in the design, implantation, and use of such devices.

We introduced the idea of performing a two-step optimization separating the implant geometry and insertion are optimized using purely geometric objective functions, and then multisite stimulation protocols are optimized for the best implant geometry and insertion. We showed that it is possible to perform stimulation protocol optimization using a surrogate model of neural activation trained on biophysical models. We showed that multipolar stimulation administered through intrafascicular TIMEs leads in theory to a substantial selectivity increase.

We hope to have provided enough evidence of the fact that modelling, machine learning and optimization methods should be used to accelerate the development and use of neuroprosthetic devices. We hope that our work will motivate the further development and use of minimally invasive structural and functional imaging techniques for peripheral nerves, to be able to reach the very high selectivities found in silico in the present work.

---

**Supplementary Figure 1.**
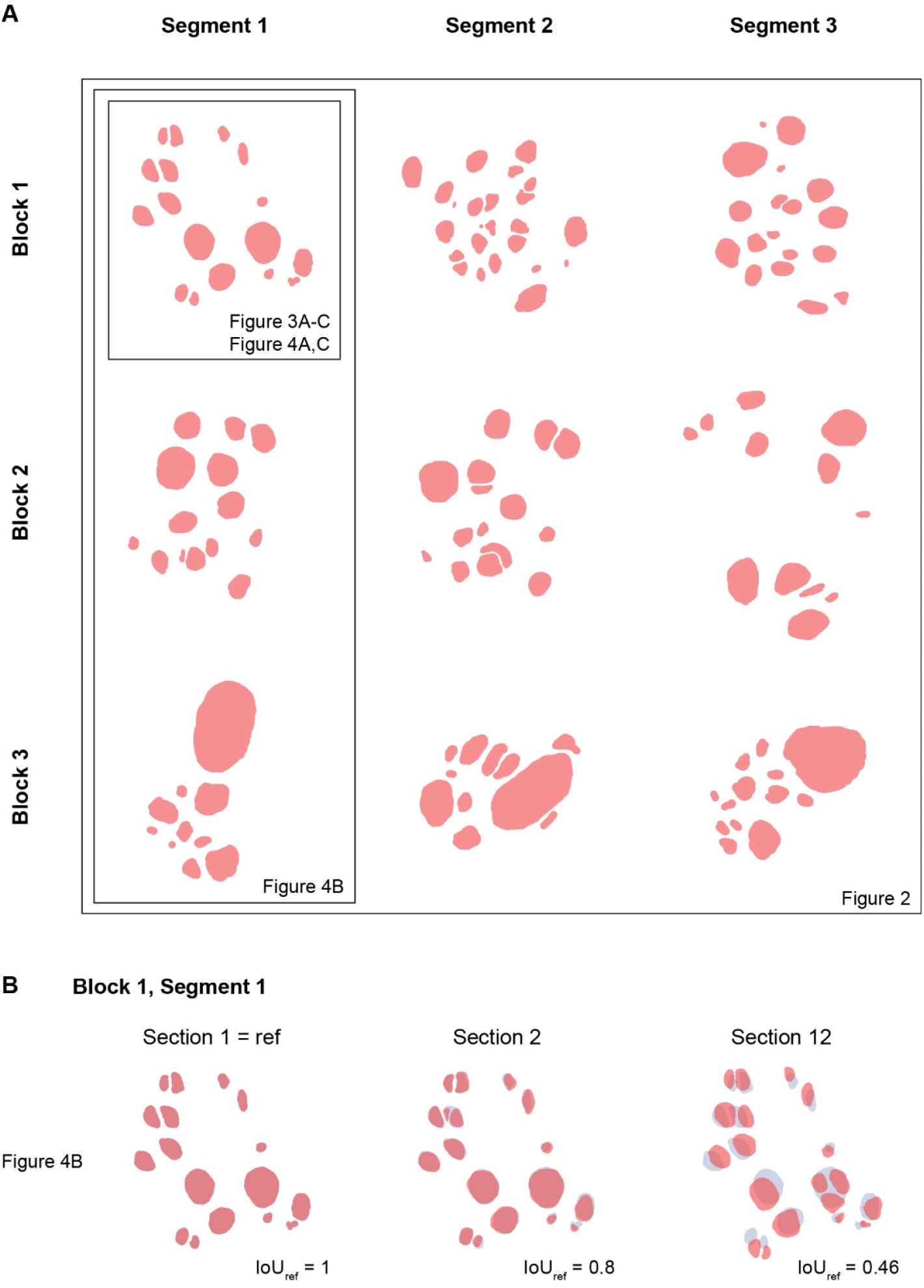
Employed nerve fascicular topographies. Nerve fascicular topographies employed in the insertion and optimization protocol optimization experiments. Pink filled polyshapes correspond to fascicles. In (B), blue filled polyshapes correspond to block 1 segment 1 section 1, and are reported for reference. IoU: intersection over union. The subsets of fascicular topographies employed in each experiment are indicated.

**Supplementary Figure 2.**
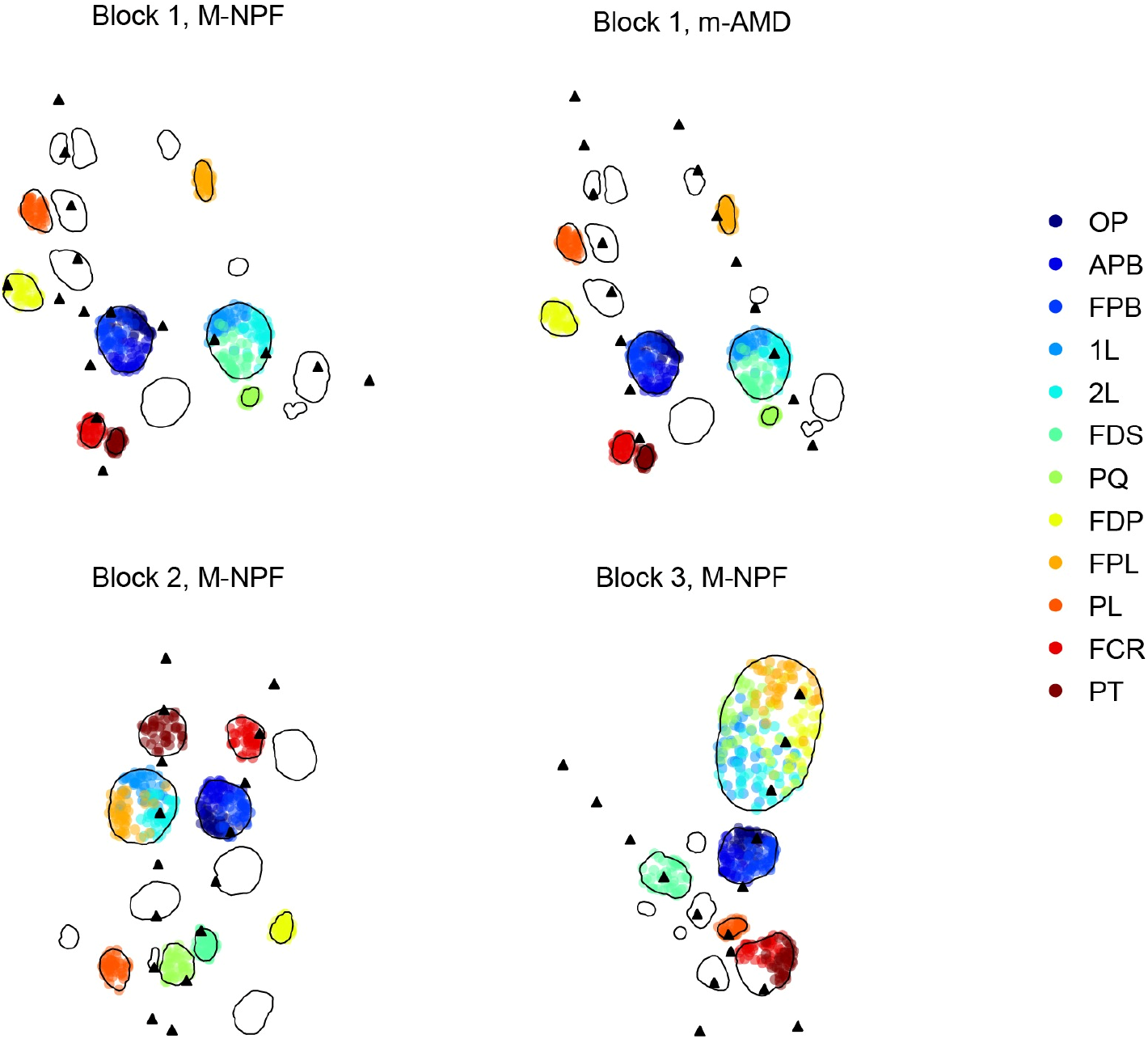
Employed nerve functional topographies. Functional topographies and implant insertions employed in the stimulation protocol optimization experiments. Solid black lines delimit nerve fascicles; colored dots correspond to nerve fibers whose color indicates the targeted muscle; black triangles correspond to stimulating site locations. Targeted muscles: opponens pollicis (OP), abductor pollicis brevis (APB), flexor pollicis brevis (FPB), first lumbrical (1L), second lumbrical (2L), flexor digitorum superficialis (FDS), pronator quadratus (PQ), flexor digitorum profundus (FDP), flexor pollicis longus (FPL), palmaris longus (PL), flexor carpi radiali (FCR), pronator teres (PT).

**Supplementary Figure 3.**
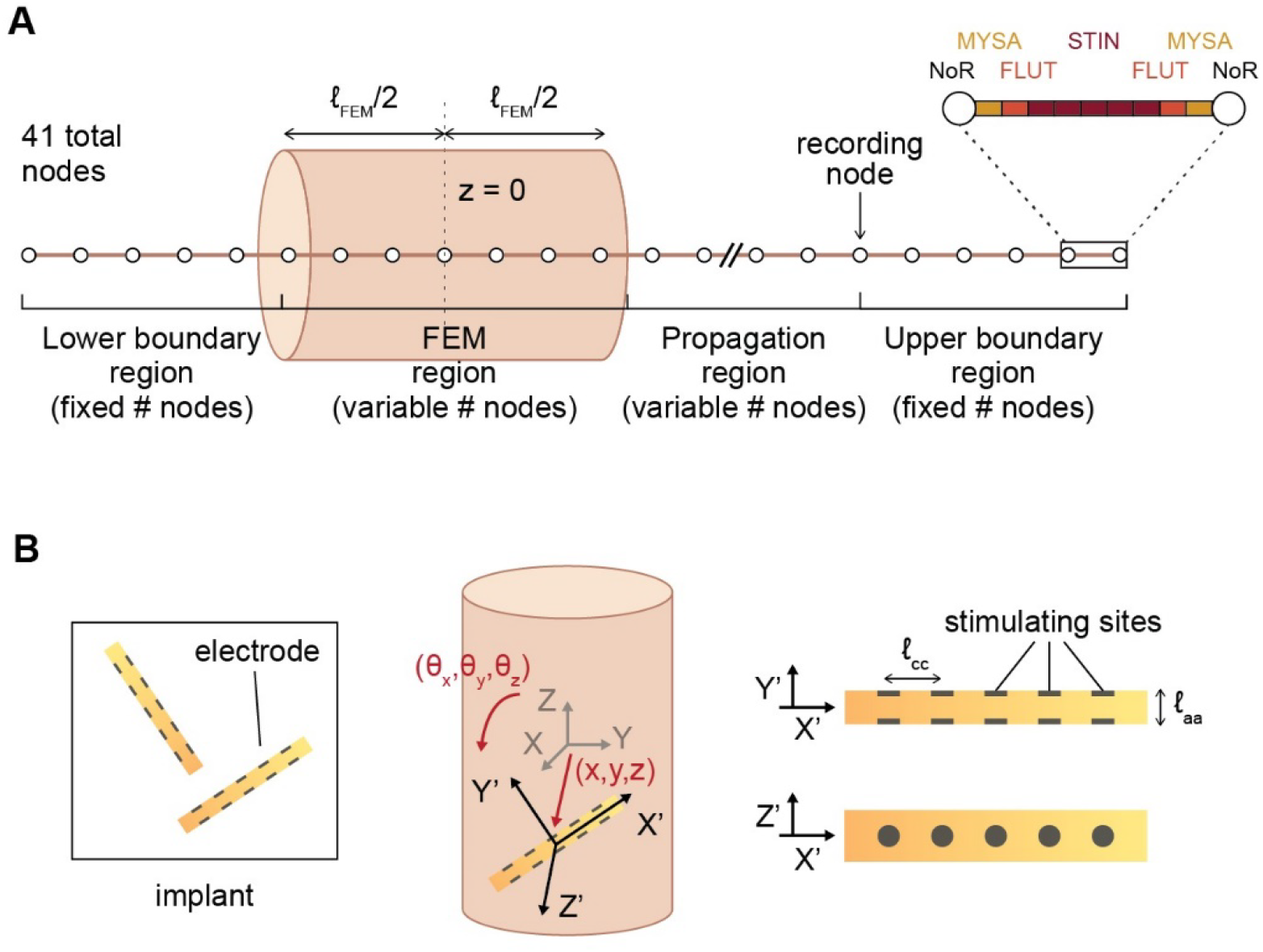
Nomenclature employed in the present work. (**A**) Subdivision and identity of fiber nodes into the different simulated regions. (**B**) Definition of implant geometry and insertion parameters. X, Y, Z: nerve / experiment reference frame; X’, Y’, Z’: electrode reference frame.

